# Potential high-frequency off-target mutagenesis induced by CRISPR/Cas9 in *Arabidopsis* and its prevention

**DOI:** 10.1101/203489

**Authors:** Qiang Zhang, Hui-Li Xing, Zhi-Ping Wang, Hai-Yan Zhang, Fang Yang, Xue-Chen Wang, Qi-Jun Chen

## Abstract

Specificity of CRISPR/Cas9 tools has been a major concern along with the reports of their successful applications. We report unexpected observations of high frequency off-target mutagenesis induced by CRISPR/Cas9 in T1 *Arabidopsis* mutants although the sgRNA was predicted to have a high specificity score. We also present evidence that the off-target effects were further exacerbated in the T2 progeny. To prevent the off-target effects, we tested and optimized two strategies in *Arabidopsis*, including introduction of a mCherry cassette for a simple and reliable isolation of Cas9-free mutants and the use of highly specific mutant SpCas9 variants. Optimization of the mCherry vectors and subsequent validation found that fusion of tRNA with the mutant rather than the original sgRNA scaffold significantly improves editing efficiency. We then examined the editing efficiency of eight high-specificity SpCas9 variants in combination with the improved tRNA-sgRNA fusion strategy. Our results suggest that highly specific SpCas9 variants require a higher level of expression than their wild-type counterpart to maintain high editing efficiency. Additionally, we demonstrate that T-DNA can be inserted into the cleavage sites of CRISPR/Cas9 targets with high frequency. Altogether, our results suggest that in plants, continuous attention should be paid to off-target effects induced by CRISPR/Cas9 in current and subsequent generations, and that the tools optimized in this report will be useful in improving genome editing efficiency and specificity in plants and other organisms.

## Introduction

CRISPR-Cas9 from microbial adaptive immune systems is a powerful tool for genome editing in a variety of organisms (Carroll and Charo, 2015; Cong et al., 2013; Jinek et al., 2012; Mali et al., 2013). In plants, various CRISPR/Cas9-based applications to basic research and crop breeding have been launched (Yin et al., 2017). For example, decades of debate on the function of *ABP1* have been resolved through CRISPR/Cas9 (Gao et al., 2015). Further analysis of the T-DNA insertion mutant of *ABP1* (*abp1-1*) found that the T-DNA insertion event also caused unwanted mutation in the adjacent *BSM* gene, thereby causing the mutant phenotype of embryonic lethality (Dai et al., 2015). Since the CRISPR/Cas9 system corrected the mistake made by the T-DNA insertion, it seemed that the CRISPR/Cas9 system was perfect. Nevertheless, the CRISPR/Cas9 system can also potentially make similar mistakes, i.e., causing undesirable mutations or off-target effects (Carroll, 2013). Thus, the case of *ABP1* not only demonstrates the huge potency of CRISPR/Cas9 tools but also provides a lesson from off-target mutagenesis to users of the tools. In addition, assessing the specificity of CRISPR/Cas9 to avoid the risk of off-target mutations, e.g., unanticipated downstream effects, is also an important regulatory concern for agricultural applications (Wolt et al., 2016).

High-frequency off-target mutagenesis induced by CRISPR/Cas9 has been reported in human cells (Fu et al., 2013; Hsu et al., 2013; O’Geen et al., 2015; Tsai and Joung, 2016; Tycko et al., 2016; Wu et al., 2014). However, it seems that off-target mutations in plants are rare and only a few cases representing low-frequency off-target mutations have been reported (Wolt et al., 2016). Whole genome sequencing (WGS) of *Arabidopsis* and rice mutants has shown that the CRISPR/Cas9 system is highly specific in plants (Feng et al., 2014; Zhang et al., 2014). Deep sequencing of a total of 178 off-target sites further demonstrated that multiplex targeting in *Arabidopsis* is highly specific to on-target sites with no detectable off-target events (Peterson et al., 2016).

Seemingly contrary to these reports, the present study reports unexpected observations of high-frequency off-target mutations induced by a sgRNA with a specificity score as high as 94 relative to the range of 0-100 (Haeussler et al., 2016; Hsu et al., 2013). Furthermore, we also describe the optimized strategies to avoid off-target mutations. Finally, we discussed the reasons for the inconsistency between this report and the previous reports.

## Results

### Unexpected observations of high-frequency off-target mutagenesis induced by CRISPR/Cas9

We previously reported the generation of 15 mutant T1 lines harboring EC1fp:Cas9 and two sgRNAs targeting the three genes: *TRY*, *CPC*, and *ETC2* (Wang et al., 2015). When we analyzed mutations in the three genes in these 15 mutant plants, we were able to obtain the *ETC2* fragment in all the 15 lines, but not able to obtain the *TRY* fragment from one line (#6) and the *CPC* fragment from 7 lines (#2, 4, 6, 8, 11, 13, and 14). After the repeated failures in the PCR amplifications, we suspected that off-target mutations might be the reason for the failures. Next, we re-conducted PCR amplifications using a modified primer pair (-FP/-RP, Figure 1) spanning the off-target and on-target sites instead of the original primer pair (-FP0/-RP, Figure 1). As a result, we successfully obtained the PCR fragments from all the lines except for the *CPC* gene in line #4, and found that the *CPC* fragments in some lines exhibited complex patterns (Figure 1). Sequencing analysis further verified that the difficulties or failures in PCR amplification as well as the complexity of the PCR products were due to off-target mutations (Table S1). Thus, we unexpectedly found high frequency (87%) off-target mutations in the *CPC* gene during analysis of the mutations in the three target genes in the 15 lines. The specificity score based on the *in silico* prediction algorithm was 94 relative to the range of 0–100 (Haeussler et al., 2016; Hsu et al., 2013), suggesting that some sgRNAs predicted to have a high specificity score may still have possibility to induce high-frequency off-target mutations. The observed differences in mutation frequencies in the three potential off-target sites, which all harbor three mismatches to the sgRNA, could be explained by previously described rules (Tsai and Joung, 2016; Tycko et al., 2016; Wu et al., 2014). The insurmountable failures in PCR amplification of the *CPC* fragment from line #4 can be attributed to T-DNA insertions, which will be described later in this report.

**Figure 1.**
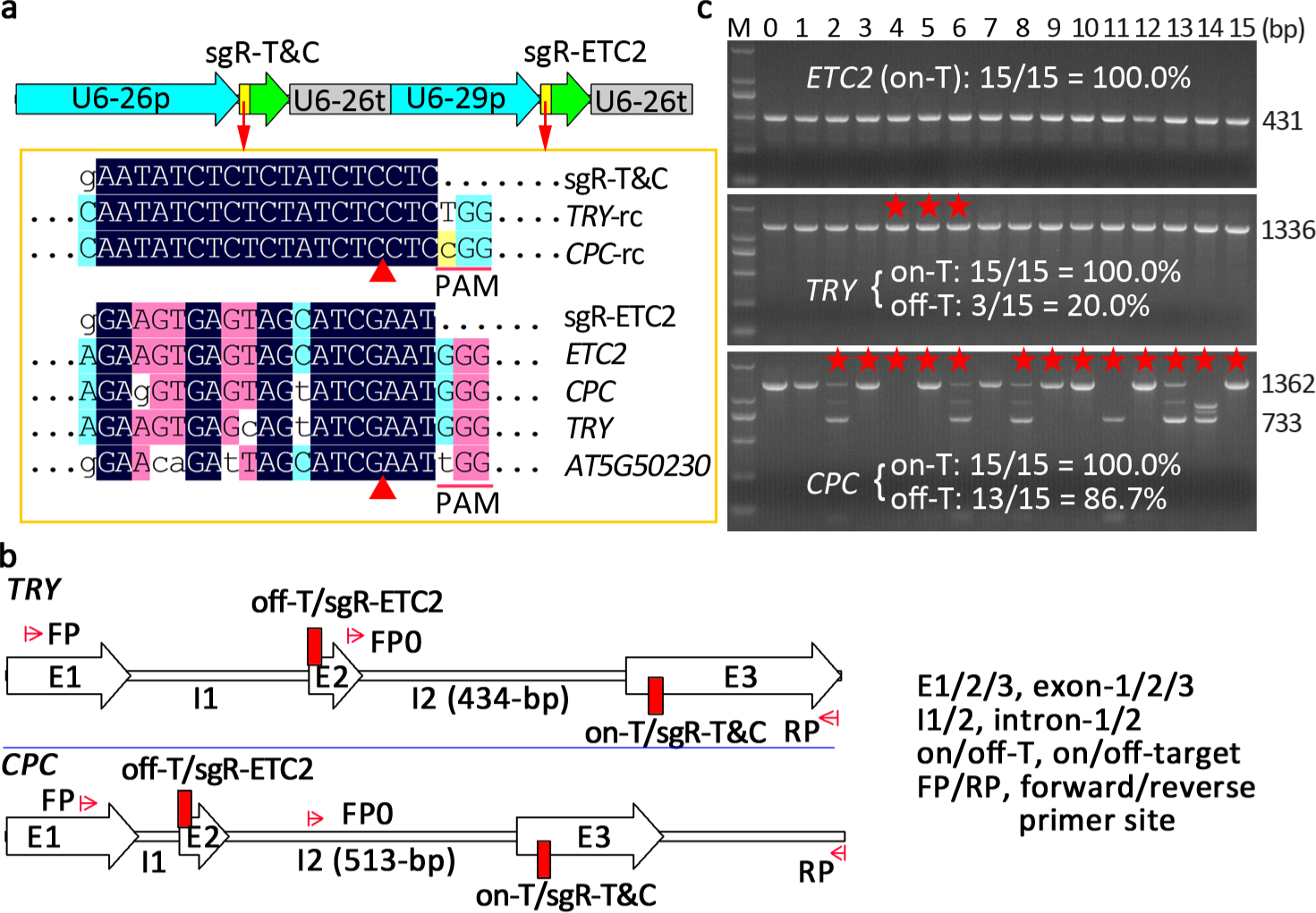
CRISPR/Cas9-induced high-frequency off-target mutagenesis in *Arabidopsis.* **(a)** Two sgRNA expression cassettes and the alignment of the sgRNAs with their target and potential off-target genes in the 15 T1 likely *try cpc etc2* triple mutant lines reported previously. Only aligned regions of interest are displayed. -rc, reverse complement. Note: the sgR-ETC2 has a specificity score of 94 relative to the range of 0-100. **(b)** The sgRNA targeting ETC2 gene (sgR-ETC2) has potential off-target sites in the two genes, *TRY* and *CPC*, which are targeted by the other sgRNA (sgR-T&C). **(c)** PCR amplification with primers spanning on-target and off-target sites revealed the sgR-ETC2-induced off-target mutations in the TRY and *CPC* genes. The stars represent off-target mutations as determined by sequencing analysis. Note: Failure in PCR amplification of the *CPC* gene in line #4 was due to T-DNA insertions described later in this report.

### Evidence of aggravated off-target effects in T2 progeny

To analyze off-target mutations in T2 plants and to simplify the analysis, we first generated two new CRISPR/Cas9 vectors, each harboring only one-sgRNA cassette. Analysis of off-target mutations indicated that the frequency of off-target mutations in the newly generated T1 plants was not as high as that in the 15 lines harboring two sgRNA cassettes, which suggests that the off-target mutations were enhanced by adjacent on-target mutations. Nevertheless, the efficiencies of the off-target mutations in the *CPC* gene of the newly generated transgenic plants were still higher than 10% (Figure 2). In practice, upon detection of unwanted off-target mutations, researchers would always pay attention to those from intended mutants rather than normal plants. Therefore, we analyzed off-target mutations in T2 plants from six *etc2* mutant lines (Figure 2). The results indicated that off-target mutations in the *CPC* gene of T2 plants were highly frequent and averaged higher than 60%. Moreover, 3% of the T2 plants harbored off-target mutations in the two genes (*CPC* and *TRY*) (Figure 2; Table S2). These results suggest that off-target effects were indeed aggravated in the T2 progeny.

**Figure 2.**
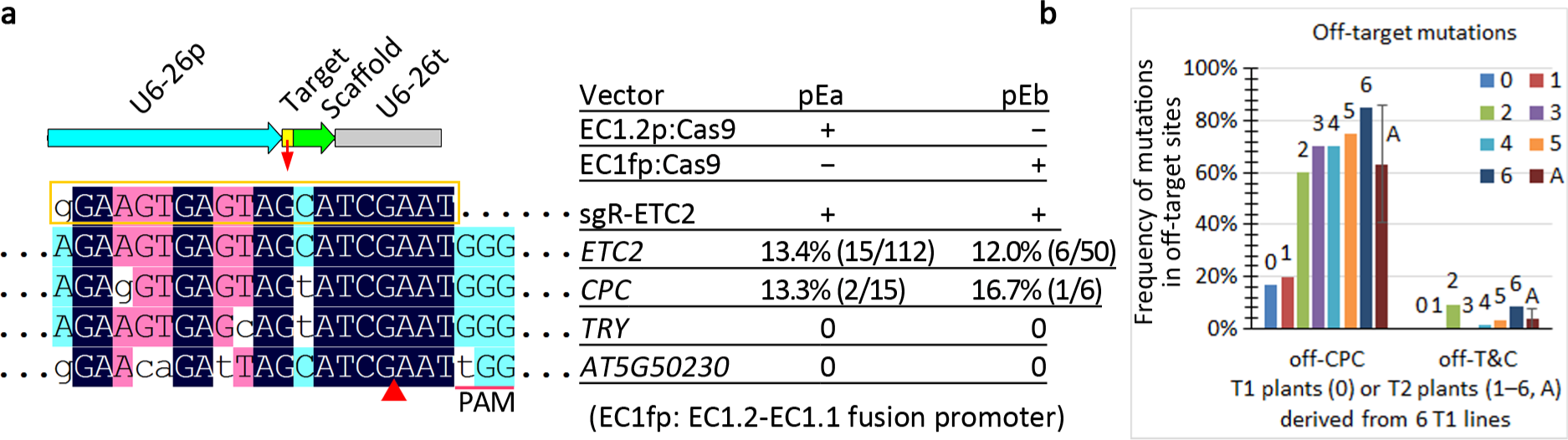
Evidence of aggravated off-target effects in T2 progeny. **(a)** On-target and off-target mutation frequencies in T1 transgenic lines harboring a sgRNA expression cassette and one of the two types of Cas9 expression cassettes. The names of the two CRISPR/Cas9 binary vectors and their Cas9/sgRNA expression cassettes are indicated on the upper right. The aligned sequences of the sgRNA, target and off-target sites are displayed on the left panel. The mutation frequencies are indicated on the right of the on/off-target genes. The on-target mutation efficiency was calculated based on the ratio of number of mutants to total number of T1 plants. The off-target mutation frequency was calculated based on the ratio of number of the mutants harboring off-target mutations to total number of mutant plants. **(b)** Off-target effects were aggravated in T2 plants. 6 T2 populations from the 6 T1 pEb transgenic plants harboring *ETC2* mutations were analyzed. For each T2 population, 20 randomly selected T2 plants were analyzed for off-target mutations in the *CPC* gene. The frequency of off-target mutations in two genes (*CPC* and *TRY*) was calculated based on the ratio of the number of mutant plants with clustered leaf trichomes to the total number of T2 plants examined. T1 line numbers 1-6 correspond to original T1 line numbers 48, 6, 31, 20, 7, and 43, of which only #20 harbors chimeric off-target mutations in the *CPC* gene in T1 plants. A, average value for the six lines.

### Optimization of the strategy for isolating Cas9-free mutants to overcome off-target mutations in the progeny

To overcome the intensified effects of off-target mutations on the T2 progeny, we introduced a mCherry cassette into our CRISPR/Cas9 system for simple and reliable isolation of Cas9-free *Arabidopsis* mutants (Gao et al., 2016). Introduction of the mCherry vector harboring the original sgRNA scaffold (Figure 3) lowered editing efficiency (2.4%) relative to that observed in our previous study (> 10% for the three target genes), suggesting that the mCherry cassette has a negative effect on the CRISPR/Cas9 system. We then made efforts to optimize the system. We used a mutant sgRNA scaffold (Dang et al., 2015), which increased editing efficiency from 2.4% to 3.8%, whereas deletion of the 3×Flag of Cas9 decreased efficiency to 1.7%. Upon fusion of the mutant sgRNA scaffold to tRNA, a significant increase in editing efficiency was observed (Figure 3). Thus, the combination of the two strategies based on the tRNA and mutant sgRNA scaffold can overcome the negative effects of the mCherry cassette on the CRISPR/Cas9 system.

**Figure 3.**
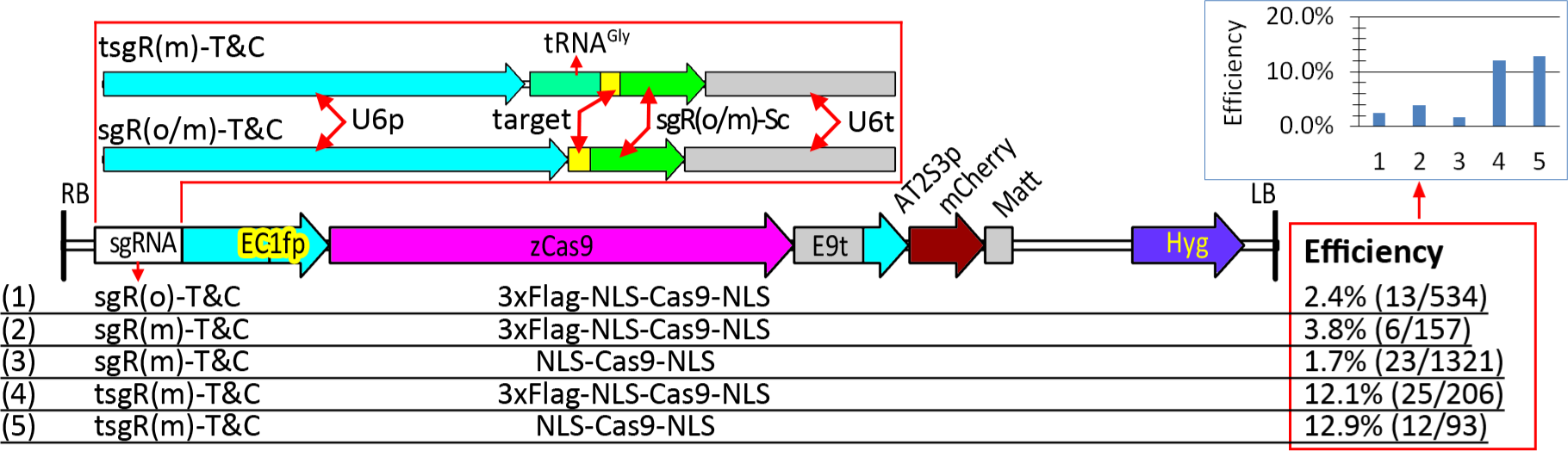
The negative effect of the mCherry cassette on editing efficiency could be overcome by fusion of tRNA with mutant sgRNA scaffold. Five combinations of three types of sgRNA cassettes (targeting *TRY* and *CPC)* and two Cas9 variants with or without 3×Flag, their physical maps of T-DNA, and efficiencies of mutations in the *TRY* and *CPC* genes of T1 plants are indicated. All the sgRNA target sequences in the five constructs were the same, and alignment of the target sequence with its target gene (*TRY* and *CPC*) is indicated in Figure 1 (sgR-T&C).

### Fusion of tRNA with mutant sgRNA scaffold conferred a significantly higher editing efficiency than that using the original sgRNA scaffold

The activities of the high-specificity SpCas9 variants have been shown to be highly sensitive to mismatches of sgRNAs to their targets; even when the first base of the sgRNAs were mismatched to their targets, editing efficiency of high-specificity SpCas9 significantly decreased (Kleinstiver et al., 2016; Slaymaker et al., 2016). Because the fused tRNA-sgRNAs were efficiently and precisely processed into sgRNAs with the desired 5' targeting sequences *in vivo* (Xie et al., 2015), the tRNA-sgRNA fusion strategy effectively facilitates the expression of the sgRNAs fully matched to their targets, thus ensuring the activities of high-specificity Cas9. To more extensively evaluate the tRNA-sgRNA fusion strategy, we generated seven constructs harboring EC1fp:Cas9 and one of the seven types of double-sgRNA cassettes (Figure 4). When we fused tRNA-Met instead of tRNA-Gly to the sgRNA targeting the *TYR* and *CPC* genes, we obtained higher editing efficiencies (Figure 4): 3.7% (pE-T&C) *vs*. 2.1% (pT&C-E), 4.2% (pE-T&C2) *vs*. 2.8% (pT&C-E2), and 2.8% (pT&C-E2) *vs*. 2.3% (pT&C-E2b). These results suggest that tRNA-Met is efficient and could be used together with tRNA-Gly for multiplex genome editing. The results also verified that the sgRNAs fully matched to targets conferred slightly higher editing efficiencies than the sgRNAs with the first base mismatched to the targets: 2.8% (pT&C-E2) *vs*. 2.1% (pT&C-E) and 4.2% (pE-T&C2) *vs*. 3.7% (pE-T&C).

**Figure 4.**
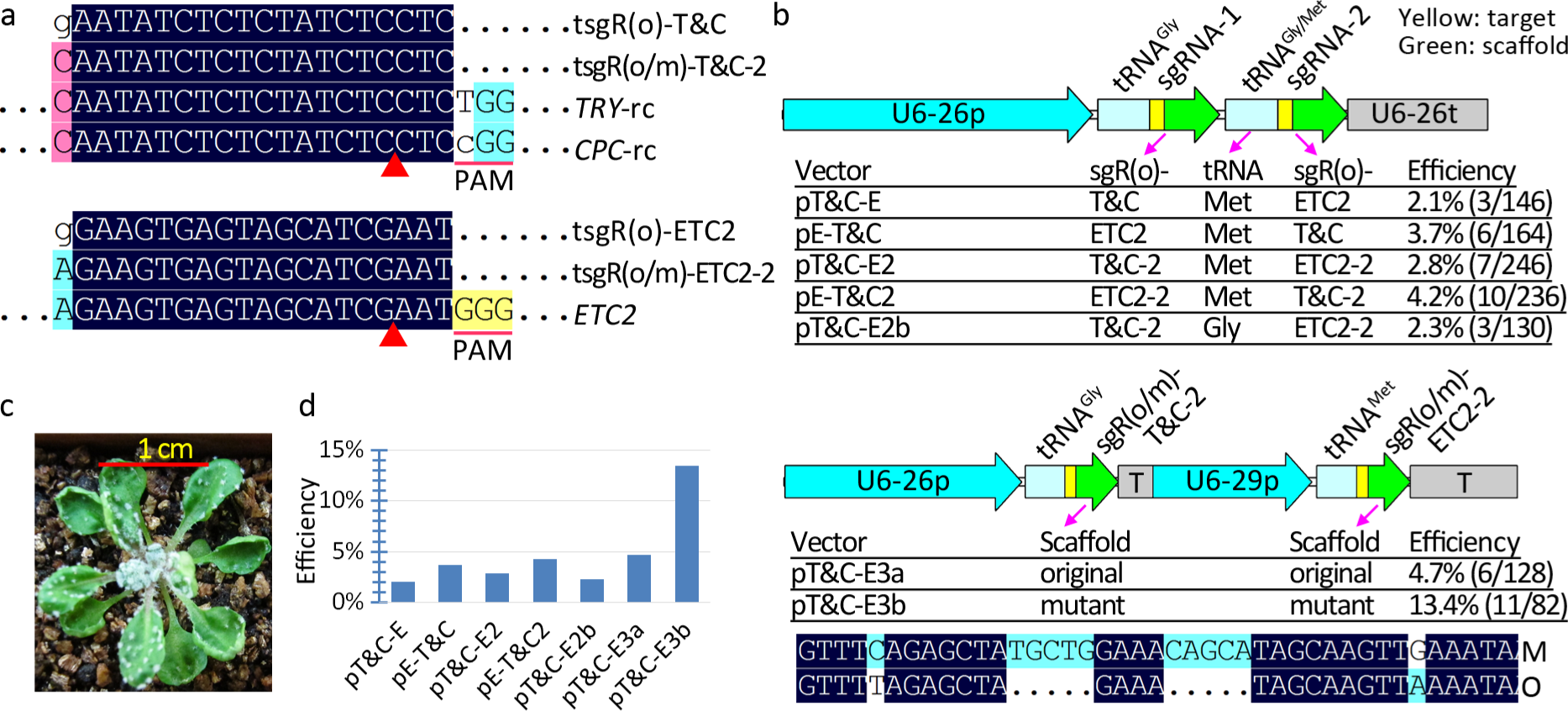
Fusion of tRNA with mutant sgRNA scaffold conferred significantly higher efficiency than with the original sgRNA scaffold. **(a)** Alignment of the sgRNAs with their target genes. Only aligned regions of interest are displayed. -rc, reverse complement. **(b)** Two strategies were used to express two tRNA-sgRNA cassettes, one using a Pol3 promoter and another using two Pol3 promoters to drive two fused tRNA-sgRNA cassettes. Seven combinations of two tRNA genes and two sgRNA scaffolds were used in testing mutation efficiencies based on the two strategies. The alignment of the original (O) and mutant sgRNA (M) scaffolds are indicated and only regions with differences are displayed. Mutation efficiencies were calculated based on the ratio of the number of mutant plants with clustered leaf trichomes to the total number of T1 plants. © Representative phenotypes of mutants with clustered leaf trichomes. **(d)** Graphic comparison of mutation efficiencies induced by the seven constructs.

Nevertheless, the above tRNA-sgRNA fusion strategies conferred significantly lower efficiencies than the general strategy we previously reported (> 10%). We postulated that two tandem tRNA-sgRNA cassettes affected the expression of sgRNA. Therefore, we used two polymerase III promoters to drive the two tRNA-sgRNA cassettes, respectively. The results indicated that this strategy was more efficient than the former: 4.7% (pT&C-E3a) *vs*. 2.8% (pT&C-E2). We then used the mutant sgRNA scaffold to replace the original one and obtained a significantly higher editing efficiency (13.4%). To further provide evidence for the efficacy of the tRNA-sgRNA fusion strategies, we first generated two vectors, each harboring one tRNA-sgRNA targeting the *ETC2* gene (Figure S1). The results proved that upon fusion to tRNA, the mutant sgRNA scaffold conferred higher editing efficiency than the original [9.2% (pEd) *vs*. 5.2% (pEc)]. The results also suggested that the off-target effects would be aggravated when the concentrations of sgRNA were significantly increased or when the mismatched base was changed to a matched one, regardless of whether the mismatch involved the first base (Figure S1). Second, we generated four additional vectors, each harboring one of the two Cas9 cassettes and one of the three sgRNA cassettes targeting the *BRI1* gene. The results similarly indicated that tRNA-sgR(m) conferred a significantly higher editing efficiency than tRNA-sgR(o), where sgR(m/o) represents the sgRNA with the mutant or original scaffold, respectively, showing 12.4% (pBd) *vs*. 1.3% (pBc) for the T1 mutant plants with observable phenotypes (Figure S2).

In summary, our results suggest that depending on the target sites, the editing efficiencies of different tRNA-sgRNA fusion strategies could be largely described in decreasing order as follows: tRNA(Met)-sgRm ≥ tRNA(Gly)-sgR(m) ≥ sgR(m) ≥ sgR(o) ≥ tRNA-sgR(o), where the sgR(m/o) represents sgRNAs with a mutant or original scaffold, respectively.

### Validation of eight high-specificity SpCas9 variants

High-specificity SpCas9 variants were developed based on two different strategies, namely, introducing mutations to weaken Cas9 binding to either the non-complementary or complementary DNA strand. These two strategies increase the stringency of guide RNA-DNA complementation for nuclease activation and therefore significantly improve editing specificity (Kleinstiver et al., 2016; Slaymaker et al., 2016). We generated eight SpCas9 variants of the maize codon-optimized Cas9 using single or combinatory forms of the two strategies (Table 1). To evaluate the editing efficiencies of the eight Cas9 variants, we used the EC1.2-EC1.1 fusion promoter (EC1fp) to drive the Cas9 variants and the tRNA-sgR(m) strategy for targeting.

**Table 1.**
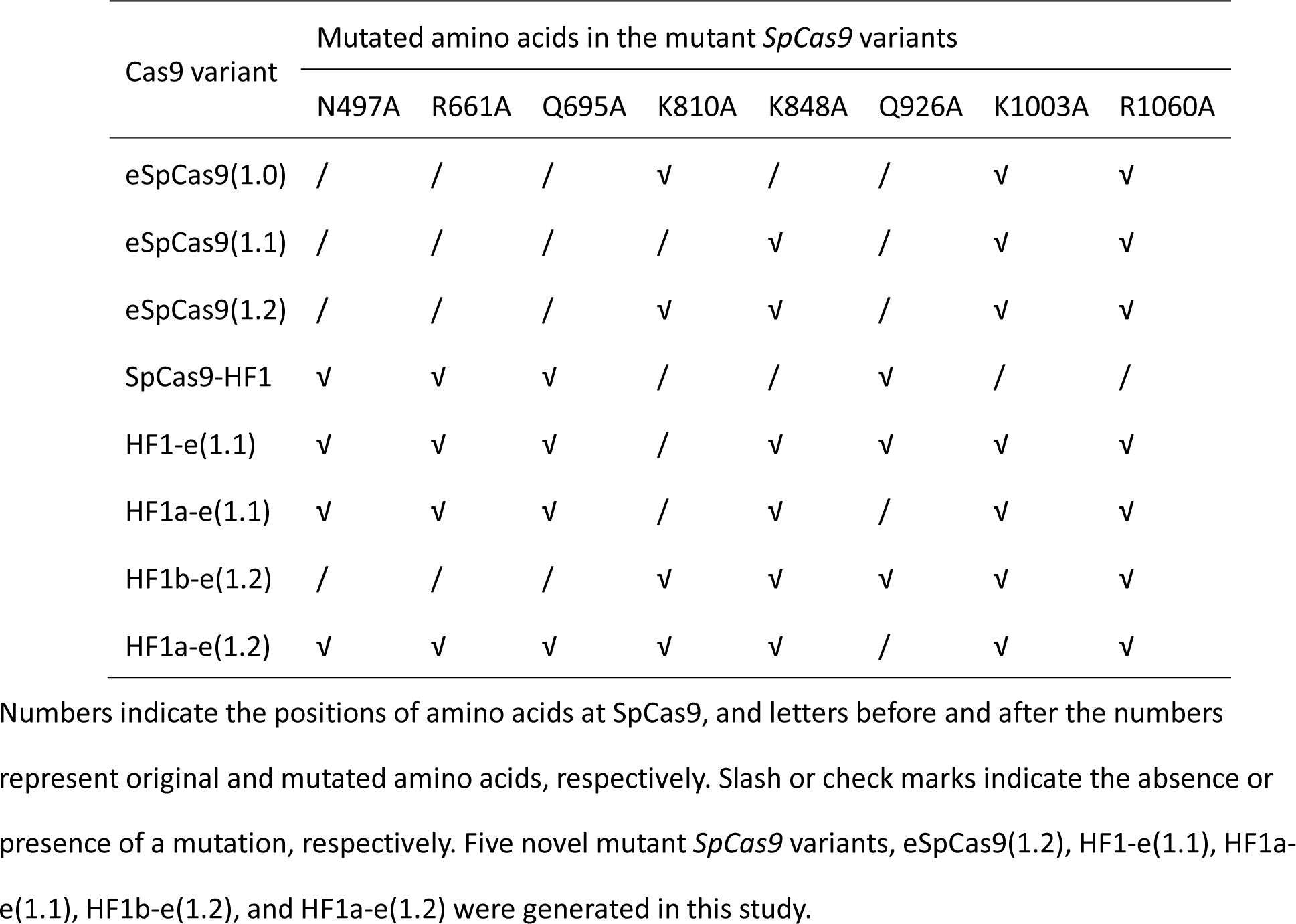
Mutant *SpCas9* variants with enhanced specificity.

We first validated the editing efficiency of two reporter genes, *CPC* and *TRY*, which exhibit observable clustered leaf trichomes when simultaneously mutated. The results indicated that only 1 of the 8 Cas9 variants displayed mutations in the T1 plants, with an editing efficiency of 0.64% (1/156), significantly lower than that (17.3%, 27/156) of the wild-type SpCas9 (Table S3). We then tested the editing efficiency of the T2 plants. The results demonstrated that the four SpCas9 variants, which were generated using combinations of mutations in two types of high-specificity SpCas9 variants, still conferred no observable mutations in the T2 plants (Table S3). The results also indicated that SpCas9-HF1 conferred significantly lower editing efficiency than eSpCas9 variants, of which eSpCas9(1.1) exhibited the highest editing efficiency (Figure 5). Interestingly, eSpCas9(1.2), the combinatory forms of eSpCas9(1.0)/(1.1), conferred high editing efficiency that was comparable to that in eSpCas9(1.0) (Figure 5).

**Figure 5.**
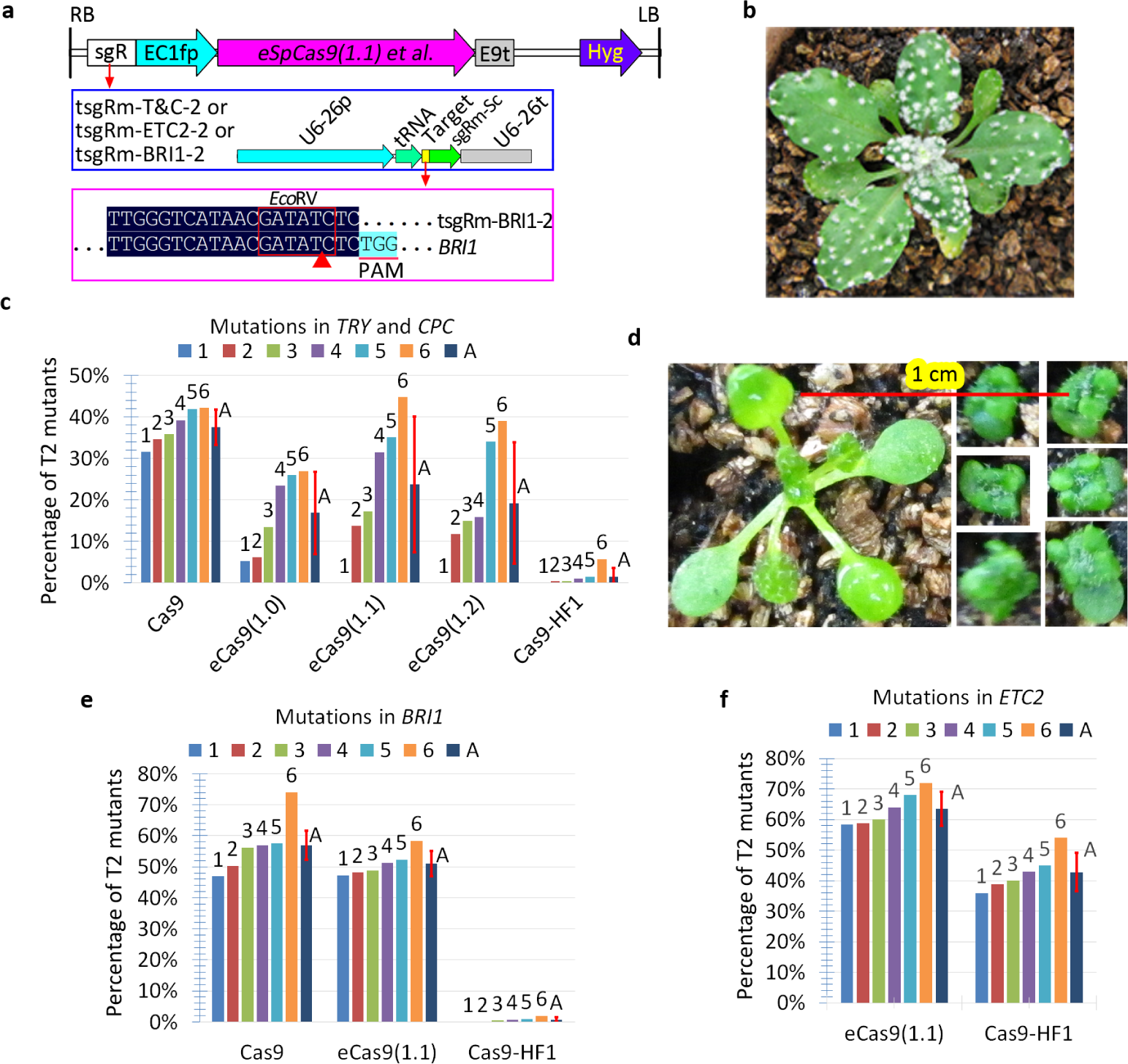
Comparison of efficiencies of mutant SpCas9 variants with high specificity. **(a)** Structures of T-DNA and the sgRNA expression cassettes for testing the efficiencies of *SpCas9* variants. Alignment of one sgRNA with its target gene *BRI1* is indicated, whereas the alignment of the other two sgRNAs with their target genes was described elsewhere. Only aligned regions of interest are displayed. **(b)** Phenotypes of a representative T2 mutant plant harboring tsgRm-T&C-2 and eSpCas9(1.1). **(c)** Efficiencies of mutations in the *TRY* and *CPC* genes of T2 plants, each harboring tsgRm-T&C-2 and one of five *SpCas9* variants. Six T1 lines (#1-6) with normal phenotypes were randomly selected for the analysis of mutations in the *TRY* and *CPC* genes of the T2 population according to clustered leaf trichomes. A, average value of the six lines. **(d)** Phenotypes of representative *bri1* mutants. One T2 plant with normal phenotypes and six segregated *bri1* mutants are displayed. The T2 transgenic plants harboring *eSpCas9(1.1)* were from the same pot and the same photo. **(e)** Efficiencies of mutations in *BRI1* of the T2 plants harboring tsgR-BRI1-2 and one of three *SpCas9* variants. Six T1 lines (#1-6) with normal phenotypes were randomly selected for analysis of mutations in *BRI1* of their T2 populations according to the dwarf phenotypes. A, average value of the six lines. **(f)** Efficiencies of mutations in *ETC2* of the T2 plants, each harboring tsgR-ETC2-2 and one of the two mutant *SpCas9* variants. Six T1 lines (#1-6) without mutations were randomly selected for analysis of mutations in *ETC2* gene of their T2 population by direct sequencing of PCR products. A, average value of the six lines.

To further compare the editing efficiencies of eSpCas9(1.1) and SpCas9-HF1, we tested two additional target genes, namely, *BRI1* and *ETC2* (Figure 5). No mutations in *BRI1* of T1 plants harboring eSpCas9(1.1) or SpCas9-HF1 were observed (Table S4). We identified eSpCas9(1.1)-induced mutations in the *ETC2* gene of the T1 plants, with an efficiency of 2.4% (1/42), significantly lower than that (9.2%, 9/98) of the wild-type SpCas9 (Table S5). For T2 plants, eSpCas9(1.1) conferred significantly higher editing efficiency of mutations in *BRI1* or *ETC2* than SpCas9-HF1. Finally, no off-target mutations induced by eSpCas9(1.1) were detected, indicating that eSpCas9(1.1) was indeed highly specific (Table S6).

Overall, our results indicated that depending on the target sites, the editing efficiencies in the T1 plants of the eight high-specificity SpCas9 could be largely described in decreasing order as follows: SpCas9 >> eSpCas9(1.1) >> the others, whereas those in the T2 plants could be largely described in decreasing order as follows: SpCas9 ≥ eSpCas9(1.1) > eSpCas9(1.0) ≥ eSpCas9(1.2) >> SpCas9-HF1 >> the others. These results suggest that high-specificity SpCas9 variants, particularly SpCas9-HF1, require significantly higher expression levels for their on-target editing efficiency. These results also suggest that the high-specificity SpCas9 variants driven by constitutive and strong promoters, in combination with tRNA-sgRNA(m) fusion strategy, enable highly efficient genome editing in crops. Editing efficiency could be further increased when the high specificity CRISPR/Cas9 systems are combined with geminivirus-based replicon systems (Cermak et al., 2017).

### High-frequency T-DNA insertions into cleavage sites of CRISPR/Cas9 targets

Because we were unable to obtain the *CPC* fragment of line #4 by PCR amplification, we then investigated the possibility of T-DNA insertions into the cleavage sites (Figure 6). As expected, the cleavage sites of the two *CPC* alleles were inserted by two T-DNAs, respectively (Figure 3). The juncture sequences between the upstream *CPC* of the cleavage site and the left border of the T-DNA were identical for the two alleles, whereas those between the downstream *CPC* of the cleavage site and the right border of T-DNA differed between the two alleles (Figure 6).

**Figure 6.**
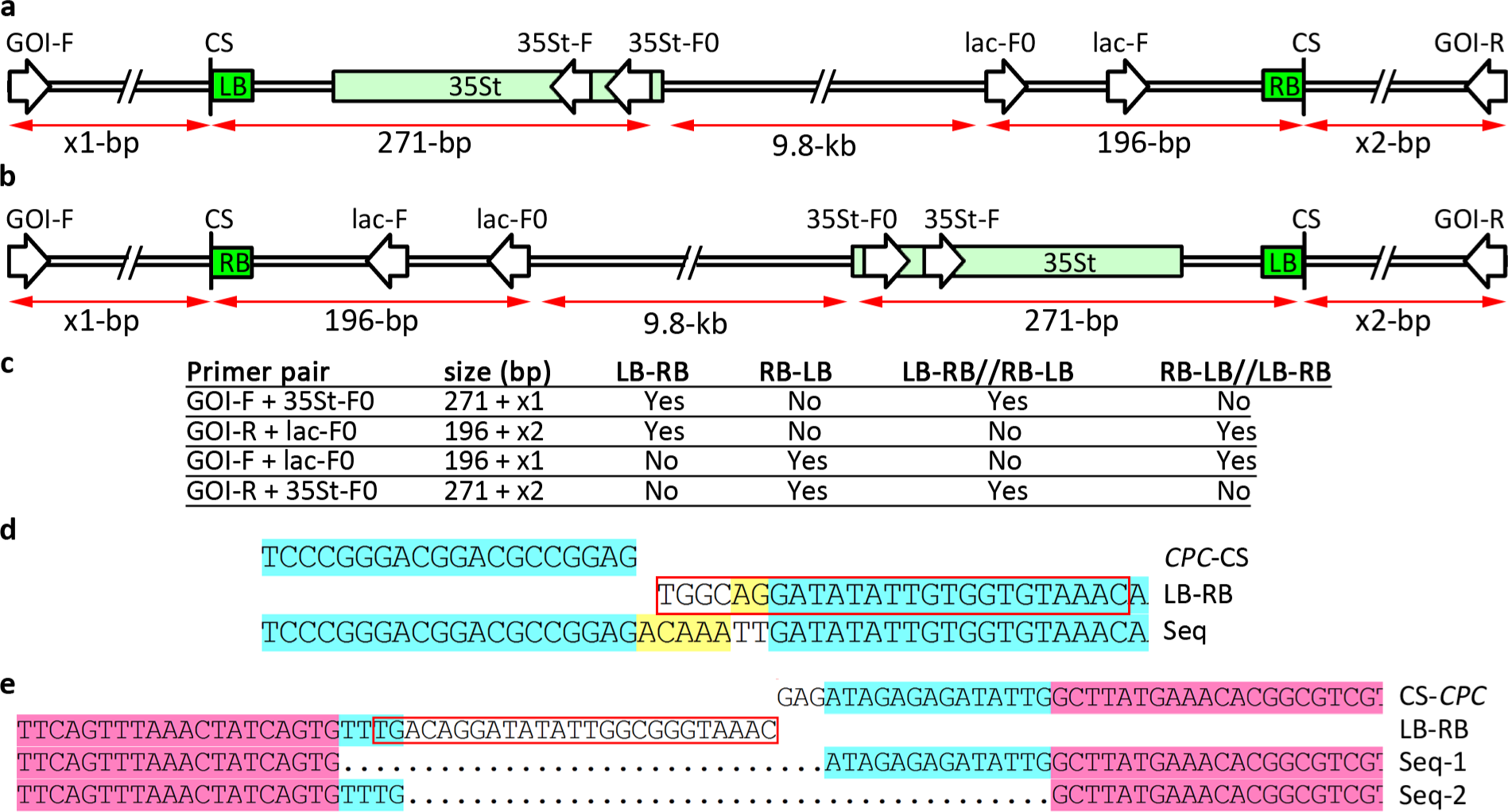
Failure in PCR amplification can be attributed to T-DNA insertions into the cleavage sites. **(a**—**c)** Schematic diagram for the identification of T-DNA insertions into a cleavage site of the CRISPR/Cas9 system. The possibly forward or reverse T-DNA insertions with one, two, or multiple copies of T-DNA, and the sizes of the PCR fragments with the indicated primer pairs are summarized. GOI, gene of interest. CS, cleavage site. -F/-R, forward/reverse primer. LB/RB, left/right border of T-DNA. 35St, CaMV 35S terminator. **(d**, **e)** Juncture sequences between the T-DNA and CPC sequences before **(d)** or behind **(e)** the cleavage site of CRISPR/Cas9 in #4 T1 lines. Note: Two types of juncture sequences between the RB and the *CPC* gene indicate that two copies of T-DNA were inserted into the two alleles, which accounted for the failure in PCR.

We also investigated 30 T1 *etc2* mutant lines generated with the three different constructs that targeted the *ETC2* gene for the possibility of T-DNA insertions into the cleavage site. The results indicated that depending on Cas9 promoters, the frequencies of targeted T-DNA insertions were 6.7% (1/15), 83.3% (5/6), and 77.8% (7/9) for the three constructs, respectively (Figure S3). In addition, we encountered the same problem for the analysis of mutations in the *HAB1* gene of line #4 T1 and T2 plants in our previous study (Zhang et al., 2017b). This failure in PCR amplifications may also be due to T-DNA insertions. As expected, we obtained juncture sequences between the downstream *HAB1* gene of the cleavage site and the T-DNA, and therefore confirmed the T-DNA insertions (Figure S4). We then analyzed the T-DNA insertions in the other 16 lines, revealing that T-DNA was inserted into the *HAB1* gene of the two additional lines, #1 and #14 (Figure S4).

SaCas9 requires a longer PAM than SpCas9 for target recognition, and therefore may be utilized in reducing off-target mutations. To validate our maize codon-optimized SaCas9, we generated a binary vector harboring EC1f:SaCas9 and U6:sgRNA. We tested a previously reported *ADH1* target (Steinert et al., 2015) and found that 15% (15/100) of the T1 plants harbored homozygous or biallelic mutations in the *ADH1* gene, indicating that the egg cell-specific promoter-controlled SaCas9 system had a similar efficiency to that of SpCas9 (Figure S5). Analysis of T-DNA insertions also indicated that 47% (7/15) of the T1 mutant lines harbored T-DNA insertions at the SaCas9 cleavage site of the *ADH1* gene (Figure S5).

## Discussion

CRISPR/Cas9 specificity is affected by various factors, including features of off-target sites and effective concentrations of the Cas9/sgRNA complexes. Determination of CRISPR/Cas9 specificity also depends on assay methods, either biased or unbiased, and *in silico* off-target prediction algorithms. This complexity is regarded as a confounding factor for off-target mutation assays and the use of different standards for measuring and reporting off-target activity affects the preciseness of conclusions (Tsai and Joung, 2016; Tycko et al., 2016; Wu et al., 2014).

CRISPR/Cas9 is known to be significantly more specific in plant cells than in human cells (Feng et al., 2014; Peterson et al., 2016; Wolt et al., 2016; Zhang et al., 2014). Two reports based on whole genome sequencing (WGS) or deep sequencing provide evidence that the editing efficiency of the CRISPR/Cas9 system is highly specific in *Arabidopsis* (Feng et al., 2014; Peterson et al., 2016). In one of these reports, deep sequencing of a total of 178 off-target sites demonstrated that the high specificity of CRISPR/Cas9 in *Arabidopsis* and the low expression levels of the Cas9 driven by the UBQ10 promoter were hypothesized to be the reason for the undetectable off-target events (Peterson et al., 2016). Actually, competitive binding of 14 sgRNA variants to Cas9 definitely led to significantly lower effective concentrations of each Cas9/sgRNA complex variant. Because out of the 14 sgRNAs, two (CLE18_2 and GLV8_1) did not show any evidence of on-target editing, the 26 off-target sites from these two sgRNAs could be excluded from the list of off-target sites. Thus, out of the rest of the 152 off-target sites from the 12 sgRNAs, 83% (126/152) and 17% (26/152) harbored 4 and 3 mismatches, respectively, indicating that the 12 sgRNAs were predicted to be specific by *in silico* analysis (Peterson et al., 2016). Therefore, lower effective concentrations of each Cas9/sgRNA complex variant and relatively specific sgRNAs might be the reasons for the reported high specificity of CRISPR/Cas9 in *Arabidopsis*.

In the other report, in-depth WGS of two T1 plants (#T1-46 and #T1-55) and one T2 plant (#T2-46) harboring GAI-sgRNA1 showed no indication of any off-target events in the potential off-target sites harboring 1-4 mismatches (Feng et al., 2014). However, although WGS is ideal for off-target mutation assays of individual plants, due to its high cost, it is not practical to systematically assess a large number of plants and sgRNA variants to determine Cas9 specificity (Wu et al., 2014). Thus, most low-frequency off target events would go unaccounted (Wu et al., 2014). Although approximately 60 T1 plants harboring GAI-sgRNA1 were examined for off-target mutations by PCR followed by sequencing, a large number of T2 plants were not examined. Because only one sgRNA and a limited number of plants from limited generations were investigated for off-target mutations, the data cannot rule out the possibility of low-frequency off-target events induced by the same sgRNA and high-frequency off-target events induced by other sgRNA variants (Feng et al., 2014).

Based on the above analysis of the results in the two reports, it is not abnormal that we observed high-frequency off-target mutations in *Arabidopsis* in our study (Figures 1 and 2), although the sgRNA was predicted to have a high specificity score (Haeussler et al., 2016; Hsu et al., 2013). Actually, the observations in this study were also inconsistent with those in our previous reports, wherein we had suggested that the same sgRNA was specific (Wang et al., 2015; Xing et al., 2014). The main reason for this inconsistency in results is that our previous conclusions were based on only one or two T1 plants, and that different promoters were used to drive Cas9. Consistent with this notion, in our previous report on sgRNA targeting the *ABI1* gene, the investigation of eight T1 *abi1* mutants and two T2 populations for off-target mutations generated similar results to that observed in the present study (Zhang et al., 2017b). It will be interesting to use our egg cell-specific promoter-controlled (EPC) CRISPR/Cas9 system to determine the specificity of GAI-sgRNAI since its specificity score was 64 relative to the range of 0-100, much lower than that (94) of the sgR-ETC2 (Haeussler et al., 2016; Hsu et al., 2013).

The high-frequency off-target mutations in the *CPC* gene could be attributed to some sequence features such as position, distribution, and identity of mismatches (Tsai and Joung, 2016; Wu et al., 2014). Although the off-target site in the *CPC* gene has three mismatches with the sgRNA-ETC2, the first mismatch located at the first base distal to PAM is usually tolerated by the CRISPR/Cas9 system (Figure 1). The second mismatch is also far away from the PAM and/or the two mismatches are situated far from each other, which may also account for the high-frequency off-target mutations in the *CPC* gene rather than the *TRY* gene or *AT5G50230* (Figure 1). The PAM-proximal ^~^11-nt were defined as the seed region for Cas9 cutting activity and mismatches in the region are less tolerated (Wu et al., 2014). In some other assays, the seed region was narrowed down to PAM-proximal 5-nt (Wu et al., 2014). Different concentrations of the CRISPR/Cas9 complex and duration of Cas9 binding and cleavage may be responsible for the observed variations in the length of the seed detected by different assays. Therefore, it is not strange for the high-frequency off-target mutations in the *CPC* off-target site harboring a mismatch in the 8^th^ nt in the PAM-proximal position (Figure 1). Similar to this study, our previous investigation involving sgRNA-ABI1 indicated that the off-target sites also harbor a mismatch in the 9^th^ nt in the PAM-proximal position (Zhang et al., 2017b). The frequency of off-target events was also affected by mismatch identity and could be largely indicated as: rN:dT ≥ rU:dG >> rC:dC >> rA/rG:dA/dG (Doench et al., 2016; Tsai and Joung, 2016; Tycko et al., 2016). The 8^th^ and 9^th^ mismatches mentioned earlier were all of the rC:dT mismatch, suggesting that this mismatch identity was frequently tolerated and contributed to the off-target effects in plants. The observation that the off-target site harboring the two mismatches to the sgRNA targeting *ABI1* displayed higher frequency off-target effects than that harboring one mismatch (Zhang et al., 2017b) indicates that more factors should be considered in the development of more precise off-target prediction algorithms that are based on large training data sets from high-throughput experiments.

Because our EPC system is a relatively short-time expression system (Wang et al., 2015), it seemed that it should have a significantly lower off-target frequency than systems driven by constitutive promoters, including 2×35S, UBQ1, UBQ10, and PcUbi4-2 (Fauser et al., 2014; Feng et al., 2014; Peterson et al., 2016). On the contrary to this supposition, we observed high-frequency off-target mutations in the *CPC* gene, which suggests that in the EPC system, a high dosage of the CRISRP/Cas9 complex in egg cells and one cell-stage embryos compensated the short duration for off-target mutations. Consistent with this notion, comparison of the results obtained with tRNA-sgRNA(m) and tRNA-sgRNA(o) suggested that the increased dosage of the complex significantly enhanced the frequency of off-target mutations: 66.7% *vs*. 20.0% (Figure S1).

It is comprehensible that tRNA-sgRNA(m) generally has a significantly higher editing efficiency than tRNA-sgRNA(o) (Xie et al., 2015) and sgRNA(m) (Dang et al., 2015). However, the present study showed that tRNA-sgRNA(o) had a markedly lower editing efficiency than sgRNA(o), quite contrary to our anticipation (Figures 3 and 4; Figures S1 and S2). This finding may be attributed to three aspects. First, the EPC system might be much more sensitive to fluctuations in effective concentrations or activities of the CRISPR/Cas9 complex than systems driven by constitutive promoters. Consistent with this notion, different terminators (Wang et al., 2015) and even the mCherry cassette behind the terminator (Figure 3) significantly affected the editing efficiency of the EPC system. In addition, high-specificity SpCas9 variants induced lower efficiency mutations in the T1 plants than their wild-type counterpart, indicating that the EPC system was more sensitive to the fluctuation in the activities of the CRISPR/Cas9 variants. For constitutively expressed CRISPR/Cas9 systems, low levels of sgRNA, if existing, from tRNA-sgRNA(o) could be compensated by the extended duration of expression or activity of the complex, thus leading to an overall high editing efficiency (Xie et al., 2015). Second, the U6 promoters we used might be more sensitive to 4×T, a potential terminator of Pol-III promoters in the original sgRNA scaffold than the OsU3 promoter. Third, the tRNA secondary structure (the cloverleaf structure) formed after transcription might enhance termination at the 4×T sites. Whatever the reason, the present study observed that the tRNA-sgRNA(m) was the optimal form that facilitated the successful application of not only the mCherry cassette in counter-selection of Cas9-free plants (Gao et al., 2016), but also high-specificity Cas9 variants (Chen et al., 2017; Kleinstiver et al., 2016; Slaymaker et al., 2016) in avoiding the occurrence of off-target effects.

Our results also suggest that high-specificity SpCas9 mutant variants require much higher concentrations to maintain high editing efficiency than the wild-type counterpart (Figure 5), although it remains to be determined whether the mutant sgRNA scaffold affected their editing efficiencies. Driven by constitutive and strong promoters in combination with the tRNA-sgRNA(m) fusion strategy, these SpCas9 variants can be used for high-specificity and high-efficiency genome editing in crops. Particularly in geminivirus-mediated CRISPR/Cas9 systems, the editing efficiency of high-specificity SpCas9 will be greatly strengthened since DNA replicons harboring Cas9 and sgRNA cassettes transiently amplify hundreds of copies in a plant cell, thus leading to very high concentrations of the CRISPR/Cas9 complex in a cell (Cermak et al., 2017). Hence, together with the rapid evolution of integrated applications of geminivirus-based replicon systems and CRISPR/Cas9 systems, high-specificity SpCas9 will be particularly useful in avoiding off-target mutations caused by high-dosage CRISPR/Cas9 complexes in a cell (Gil-Humanes et al., 2017; Wang et al., 2017).

One of our unexpected but interesting findings in this report was the high-frequency T-DNA insertions into cleavage sites (Figure 6; Figures S3-S5). This finding could facilitate the experimental design for targeted integration of transgenes (Tzfira et al., 2003) and may trigger additional concerns for mutation analysis. First, when encountering a failure in PCR amplification of a target region, T-DNA insertions should be considered as a possibility. Second, in previous reports, the mutation types that were identified as insertions of unknown large fragments might be re-considered as T-DNA insertion events. Third, for T1 *Arabidopsis* mutants generated using the EPC system or T0 mutants generated from embryogenic callus, ratios of homozygous or biallelic mutants are subject to overestimation and underestimation, respectively. The targeted integrations of two copies of T-DNA into the two alleles of the *CPC* gene in this report or the *HAB1* gene in a previous study might also represent novel knowledge of Agrobacterium-mediated T-DNA insertions into plant genome. First, CRISPR/Cas9-mediated DNA cleavage could be completed before T-DNAs are integrated into the genome of target cells. Second, for *Arabidopsis* floral dip transformation (Desfeux et al., 2000), random integration of T-DNA usually occurs before fertilization, whereas CRISPR/Cas9-mediated targeted integration of T-DNAs occurs after fertilization.

Since the sgR-ETC2 induced off-target mutations in *CPC* with high frequency, in *TRY* with medium frequency, and in *AT5G50230* with low frequency under detectable level, our results suggest that carefully selected target sites largely guarantee high specificity. This observation was supported by our previous report, wherein we detected sgR-ABI1-induced off-target mutations in *ABI2* and *AT5G02760*, but did not detect off-target mutations in *AT2G25070* and *AT3G17090*. The intended on-target sites should have no potential off-target sites that harbor less than three mismatches and are easily tolerated mismatch features. In addition, awareness is advised when targeting multiple highly homologous genes because off-target sites are possibly adjacent to on-target sites, which may lead to enhanced off-target effects, similar to the case reported in this study. Overall, we recommend using combinatory forms of the following six strategies to avoid off-target effects in plants (Tycko et al., 2016). First, high-specificity targets should be carefully selected using *in silico* predictions (Haeussler et al., 2016). Second, Cas9-free mutants should be isolated as far as possible (Gao et al., 2016; Lu et al., 2017). Third, when necessary, high-specificity SpCas9 variants in combination with the tRNA-sgRNA(m) fusion method can be used (Chen et al., 2017; Kleinstiver et al., 2016; Kulcsar et al., 2017; Slaymaker et al., 2016; Xie et al., 2015; Zhang et al., 2017a). Fourth, when necessary, SaCas9 or other orthologs with higher specificity can be used (Steinert et al., 2015). Fifth, when necessary, paired nickases can be used (Fauser et al., 2014). Last, when necessary, DNA-free methods can be employed (Liang et al., 2017; Svitashev et al., 2016; Woo et al., 2015).

Collectively, this is the first report on the observations of high-frequency off-target mutagenesis induced by CRISPR/Cas9 in plants; our results suggest that in plants harboring CRISPR/Cas9 components, continuous attention should be paid to off-target effects induced by CRISPR/Cas9 in current and subsequent generations, and that the tools optimized in this report will be useful to improve genome editing efficiency and specificity in plants and other organisms.

## Materials and Methods

### Vector construction

All primers used in this study are listed in Table S7. The cloning CRISPR/Cas9 binary vectors, the PCR template vectors, the final CRISPR/Cas9 binary vectors each harboring one sgRNA cassette, and the final CRISPR/Cas9 binary vectors each harboring two sgRNA cassettes are listed in Tables S8–11. Detailed descriptions of the vector construction and annotated sequences of the sgRNA cassettes for cloning are provided in Methods S1 and Appendix S1–S5, respectively.

### Generation of transgenic *Arabidopsis* plants and analysis of mutations and T-DNA insertions

We transformed the 26 and 7 final CRISPR/Cas9 binary vectors, harboring one and two sgRNA cassettes, respectively, into *Agrobacterium* strain GV3101. We generated transgenic plants by transformation of *Arabidopsis* Col-0 wild-type plants via the floral dip method. We screened the seeds collected from the transformed plants on MS plates supplemented with 25 mg/L hygromycin and transplanted the resistant seedlings (T1) to soil.

To analyze mutations, we extracted genomic DNA from T1 or T2 transgenic plants grown in soil. We amplified fragments surrounding on-target and adjacent off-target sites by PCR using gene-specific primers (Table S7). We submitted purified PCR products for direct sequencing with the corresponding primers. We cloned poorly sequenced PCR products and submitted individual positive clones for sequencing using the T7 and SP6 primers.

To analyze T-DNA insertions into the cleavage sites, we used the T-DNA LB/RB-specific primer 35St-F0/lac-F0 and the gene-specific primers (Table S7) for PCR amplifications of the juncture sequences between the T-DNA and target genes. We submitted the purified PCR products for direct sequencing with primers 35St-F/lac-F (Table S7). We cloned poorly sequenced PCR products and submitted individual positive clones for sequencing using the T7 primer.

## Acknowledgements

This work was supported by grants from the National Crop Breeding Fund (2016YFD0101804), the National Natural Science Foundation of China (31670371), the National Transgenic Research Project (2016ZX08009002), the Chinese Universities Scientific Fund (2017TC007), and Collaborative Innovation Center of Crop Stress Biology, Henan Province.

## Supporting information

Additional Supporting Information may be found online in the supporting information tab for this article:

**Figure S1.** Strategy of tRNA-sgRNA(m) fusion conferred much higher efficiency of mutations in the *ETC2* gene than that with tRNA-sgRNA(o) fusion.

**Figure S2.** Strategy of tRNA-sgRNA(m) fusion confers significantly higher efficiency of mutations in the *BRI1* gene than that with tRNA-sgRNA(o) fusion.

**Figure S3.** Juncture sequences between T-DNA and *ETC2* sequences before or behind the cleavage site of CRISPR/Cas9.

**Figure S4.** Juncture sequences between T-DNA and *HAB1* before or behind the cleavage site of CRISPR/Cas9.

**Figure S5.** Juncture sequences between T-DNA and *ADH1* before or behind the cleavage site of CRISPR/SaCas9.

**Table S1.** On-target and off-target mutation analysis of 15 T1 mutant plants.

**Table S2.** Frequencies of off-target mutations in T1 and T2 plants.

**Table S3.** Efficiencies of mutations in *TRY* and *CPC* induced by mutant SpCas9 variants.

**Table S4.** Efficiencies of mutations in *BRI1* induced by mutant SpCas9 variants.

**Table S5.** Efficiencies of mutations in *ETC2* induced by mutant SpCas9 variants.

**Table S6.** Frequencies of off-target mutations induced by eSpCas9(1.1).

**Table S7.** Primers used in this study.

**Table S8.** Features of cloning CRISPR/Cas9 binary vectors.

**Table S9.** Structures of the PCR templates used in the assembly of sgRNA cassettes.

**Table S10.** Final CRISPR/Cas9 binary vectors, each harboring one sgRNA cassette.

**Table S11.** Final CRISPR/Cas9 binary vectors each harboring two sgRNA cassettes.

**Methods S1.** Vector construction.

**Appendix S1–S5.** Annotated sequences of the sgRNA cassettes for cloning.

